# Run or die in the evolution of new microRNAs - Testing the Red Queen hypothesis on *de novo* new genes

**DOI:** 10.1101/345769

**Authors:** Yixin Zhao, Guang-An Lu, Hao Yang, Pei Lin, Zhongqi Liufu, Tian Tang, Jin Xu

## Abstract

The Red Queen hypothesis depicts evolution as the continual struggle to adapt. According to this hypothesis, new genes, especially those originating from non-genic sequences (i.e., *de novo* genes), are eliminated unless they evolve continually in adaptation to a changing environment. Here, we analyze two Drosophila *de novo* miRNAs that are expressed in a testis-specific manner with very high rates of evolution in their DNA sequence. We knocked out these miRNAs in two sibling species and investigated their contributions to different fitness components. We observed that the fitness contributions of miR-975 in *D. simulans* seem positive, in contrast to its neutral contributions in *D. melanogaster*, while miR-983 appears to have negative contributions in both species, as the fitness of the knockout mutant increases. As predicted by the Red Queen hypothesis, the fitness difference of these *de novo* miRNAs indicates their different fates.

## Introduction

Organisms evolve to adapt to changing environments via the acquisition of adaptive traits. Adaptations are generally transient because continual changes are needed to keep pace with a moving world, as the Red Queen advises Alice (Carroll 1893). Van Valen (1973) emphasized biotic factors as the driving selective force, which has subsequently been expanded to include the coevolution of host/parasite, prey/predator, male/female (sexual selection), as well as antagonistic evolution (Tobler and Schlupp 2005; King, et al. 2009; Brockhurst 2011; Liow, et al. 2011; Morran, et al. 2011; Brockhurst, et al. 2014; Nordbotten and Stenseth 2016; Greenspoon and Mideo 2017). Nevertheless, the essence of this metaphor is continual adaptation, which could be driven by biotic, abiotic or both factors. Van Valen (1973) first reported that genera and species, regardless of their age, often become extinct when they cannot sufficiently adapt. We proposed to test the Red Queen hypothesis by studying the evolution of new genes, which can be compared to those in younger taxa. If the adaptive landscape shifts rapidly, as posited by the Red Queen hypothesis, this may suggest that newly emerging genes have been under strong pressure to evolve continually or face elimination.

New genes fall in two broad categories: those that originated from reshuffling of existing gene components and those emerging *de novo* from non-genic sequences (Long, et al. 2013; Andersson, et al. 2015; Schlotterer 2015; VanKuren and Long 2018). The majority of new genes belong to the former, comprising duplicated, (retro-)transposed, and chimeric genes (Chen, et al. 2013). In comparison, *de novo* genes are more appropriate for testing the Red Queen hypothesis because their products are truly new in the cellular environment. The most common source of *de novo* genes may be small non-coding loci represented by microRNAs (miRNAs) (Lu, Shen, et al. 2008; Lyu, et al. 2014). Given the ease with which miRNAs can be formed *de novo* (Meunier, et al. 2013; Lyu, et al. 2014), many could have emerged adaptively (Lu, Fu, et al. 2008; Mohammed, et al. 2013; Lyu, et al. 2014; Mohammed, et al. 2014; Mohammed, et al. 2018) although the majority may be functionless (Lu, Shen, et al. 2008; Berezikov, et al. 2010), or dead-on-arrival (Petrov, et al. 1996; Petrov and Hartl 1998). Following the molecular evolutionary analysis performed by Lyu et al. (2014), we use the term “*de novo* miRNAs” to specifically refer to the first group of adaptive miRNAs. We hypothesize that “*de novo* miRNAs” must continuously evolve and survive in different fitness landscapes or face elimination, as inferred by the Red Queen hypothesis.

In order to test this possibility, experiments must be conducted in close sibling species, as their divergence time is sufficiently short such that rapidly evolving genes remain identifiable. More importantly, they provide distinct fitness landscapes, and thus *de novo* genes might end up with different fates. Therefore, we performed functional and molecular analysis on two typical *de novo* miRNA genes (miR-983 and miR-975) in closely related *Drosophila* species that diverged about 5 – 50 Myrs ago (Kumar, et al. 2017).

## Result

### Sequence and expression divergence of miR-983 and miR-975 in *Drosophila*

We collected and sequenced 14 small RNA libraries from the testes of 5 *Drosophila* species to survey the sequence and expression patterns of the *de novo* miRNAs (**Fig. 1A**). miR-975 is present in all *Drosophila* species, while miR-983 is absent in *D. virilis*. Similar to most young miRNA genes, these two *de novo* miRNAs have rapidly evolved across species, as divergence can be seen in both the seed and non-seed regions compared to conserved miRNAs (bantam and miR-184) (**Fig. 1B**). The number of substitutions in these miRNAs is far higher than can be sustained by the neutral process, suggesting adaptive evolution (Lyu, et al. 2014; Mohammed, et al. 2014). In contrast, bantam and miR-184 are highly conserved across *Drosophila* species, even in non-seed regions, a common trend among older miRNAs.

**Figure 1.**
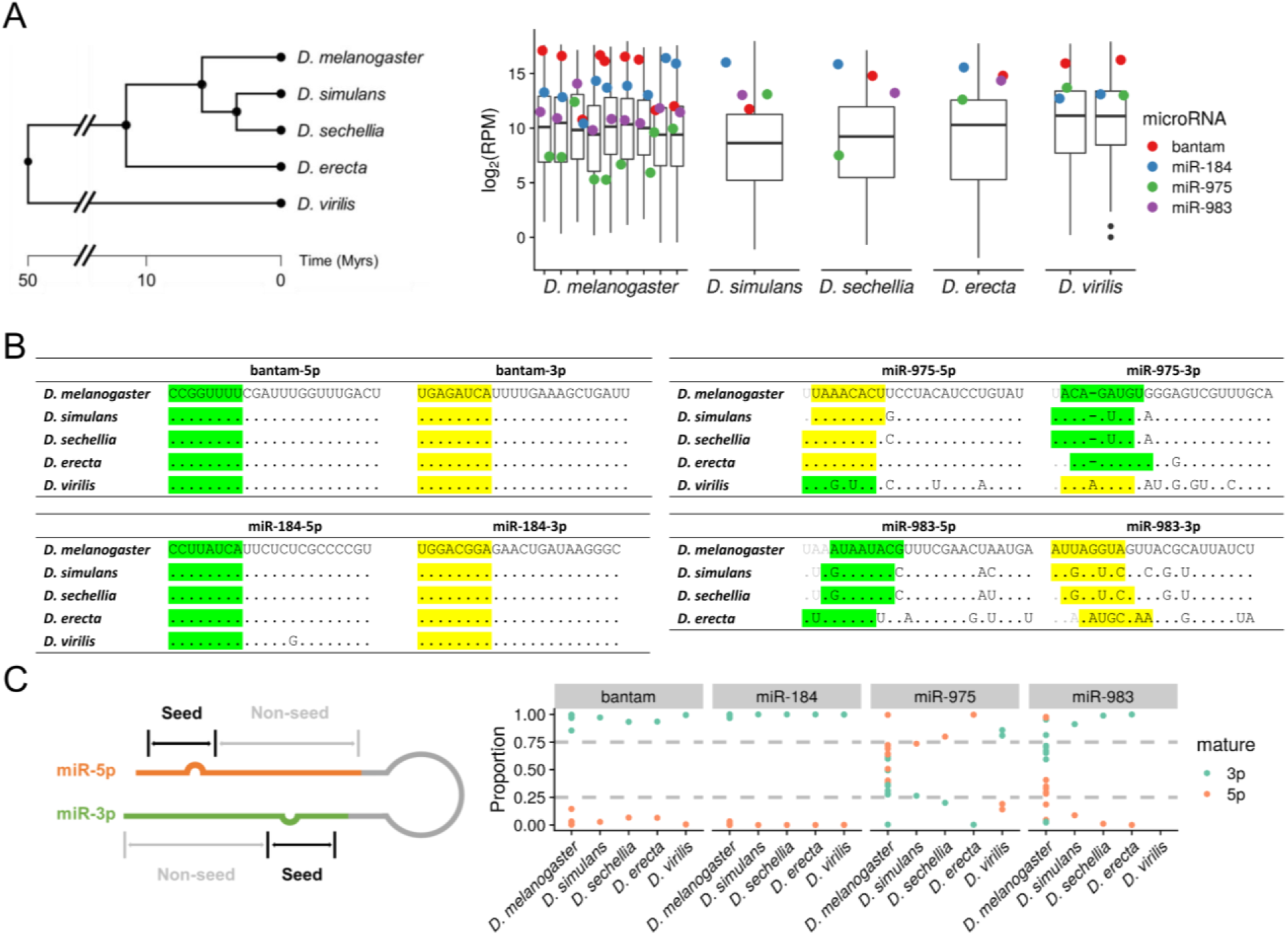
Sequence and expression divergence of miR-975 and miR-983 in Drosophila species. A) The phylogenetic tree of five Drosophila sibling species is shown on the left. Adopted from TimeTree (Kumar, et al. 2017), scale in million years (Myrs). Expression levels (Reads Per Million, RPM) of miR-975 and miR-983 are shown on the right. Boxplot represents the expression level of all detected miRNAs, and dots represent the expression of selected miRNAs annotated with different colors. miR-5p and miR-3p expression levels are summed together for comparison. B) Sequences of two highly conserved miRNAs, bantam and miR-184, across five Drosophila species are shown on the left. Sequences of miR-975 and miR-983 in the same species are depicted on the right. Seed regions of major and minor miRs are highlighted in yellow and green, respectively. C) Arm switching of miR-975 and miR-983 in different Drosophila species. Relative expression levels of miR-5p and miR-3p are shown across species (right panel), with an annotation sketch of the miRNA structure (left panel).

We then examined the expression of these miRNAs in different species and multiple lines of *D. melanogaster* and *D. virilis* as available (**Table S1**). **Figure 1A** shows the expression of bantam, miR-184, miR-975 and miR-983 relative to all miRNAs detected in the testes. Bantam and miR-184 are highly expressed and almost always ranked in the first quantile among the evaluated *Drosophila* species. In contrast, miR-975 shows substantial variation between and even within species. In *D. simluans*, *D. erecta* and *D. virillis*, its expression is high and comparable with bantam and miR-184, while in *D. sechellia*, its expression drops below the average level. In *D. melanogaster*, miR-975 expression is generally low, ranking in or close to the fourth quantile in six of nine samples. Compared with miR-975, miR-983 shows less variation across species and has higher expression, except in *D. simulans*.

Each miRNA locus, either new or old, can potentially generate two mature products, which are usually referred as the −5p and −3p arm, respectively (Ameres and Zamore 2013). The starting position of each arm may shift, resulting in distinct mature miRNAs. For example, miR-983-3p has four distinct seeds in four sibling species, and miR-975-5p has three distinct seeds in five sibling species (**Fig. 1B**), which determines the repertoire of target genes and, presumably, their functions. We also observed arm switching for young miRNAs among different species (Berezikov 2011; Zhao, et al. 2018; Kim, et al. 2020). **Figure 1C** shows that the conserved miRNAs (bantam and miR-184) predominantly express one product. In contrast, the expression levels of miR-983 and miR-975 are divergent among *Drosophila* species. For example, expression of miR-975-3p is strong in *D. virilis* but rather weak in *D. simluans, D. sechellia* and *D. erecta* (**Fig. 1C**). These three sibling species show arm-switching vis-à-vis *D. virilis,* with the 5p miR being the major form. We should also note that the divergence pattern can be observed even between different lines of the same *D. melanogaster* species.

In short, the mature products of *de novo* miRNAs are extremely variable in sequence, expression and biogenesis among species, which indicates their rapid evolution across different species.

### Rapid functional evolution of miR-983 and miR-975

We then surveyed the functional evolution of miR-983 and miR-975 by constructing loss-of-function mutants. For a direct comparison with our previous constructs in *D. melanogaster* (Lu, et al. 2018), we used TALEN (Transcription activator-like effector nuclease) to construct miRNA mutants in *D. simulans*, a non-model organism (**Fig. S1**). Mutants were confirmed by PCR followed by DNA sequencing (**Fig. 2A**), and the absence of mature miRNA expression was further validated by RT-qPCR in testes (**Fig. S2**).

**Figure 2.**
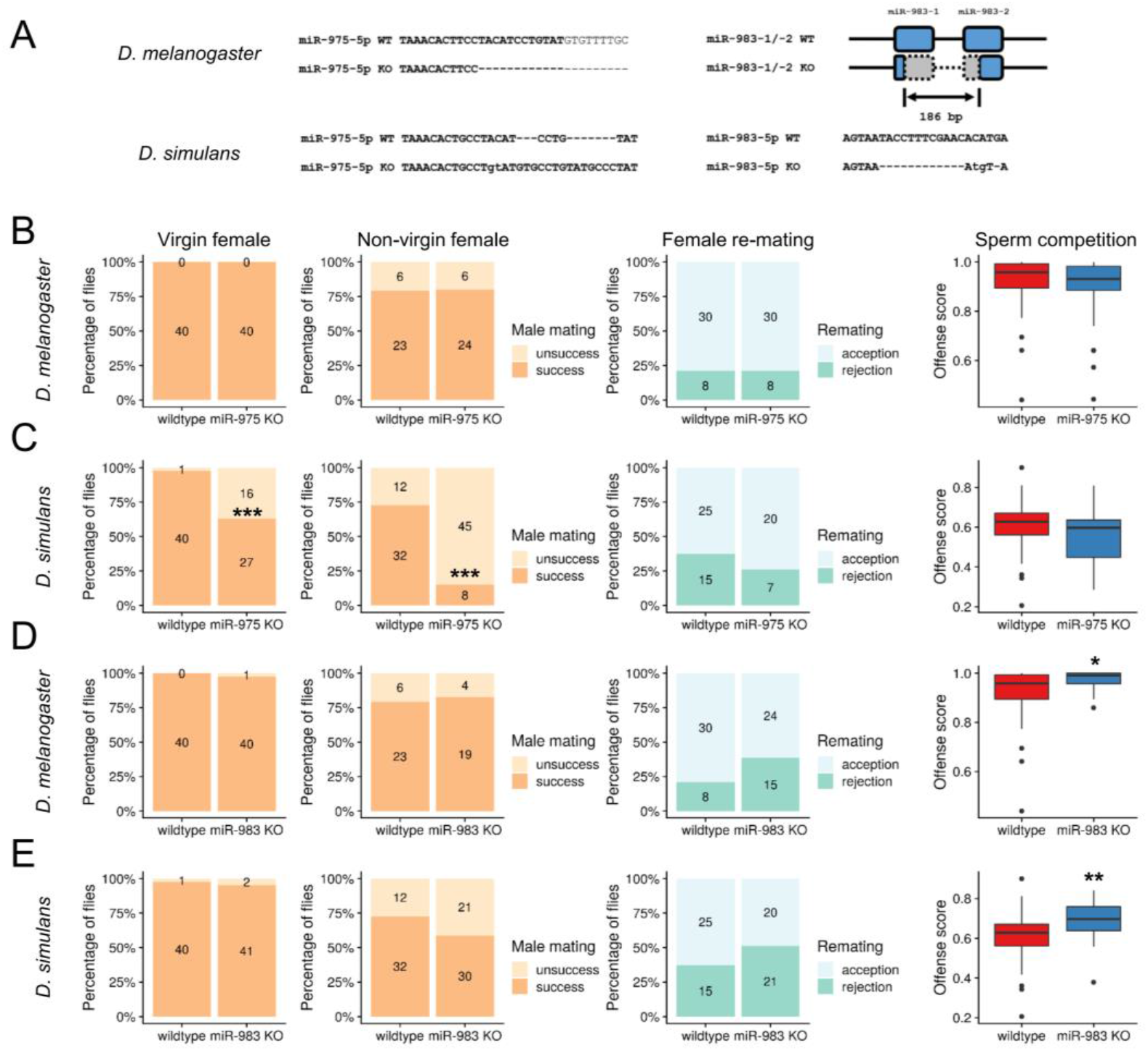
Functional divergence of miR-975 and miR-983 in *D. melanogaster* and *D. simulans*. A) miR-975 and miR-983 mutant sequences in *D. melanogaster* and *D. simulans*. miR-983 has two sequence identical paralogs in *D. melanogaster*: *dme-mir-983-1* and *dme-mir-983-2*. A 186-bp deletion disrupts both copies. WT, wildtype. KO, knockout. B) Phenotypic assays for miR-975 KO in *D. melanogaster*. Four fitness measurement are shown from left to right: male mating success with virgin and non-virgin females, Fisher’s exact test was used for significance, *** represents p ≤ 0.001; male ability to repress female re-mating; offense score for sperm competition, Mann-Whitney U test was used for significant test, * represents p ≤ 0.05 and ** represents p ≤ 0.01. C) Phenotypic assays for miR-975 KO in *D. simulans* D) Phenotypic assays for miR-983 KO in *D. melanogaster*. E) Phenotypic assays for miR-983 KO in *D. simulans* Phenotypic assays for miR-975 and miR-983 in *D. melanogaster* are adopted from Lu et al. (2018).

These KO lines were then subjected to a series of analyses focusing on the typical fitness components associated with male reproductive functions, including male mating success, male ability to repress female re-mating, sperm competition, and male fertility. Male mating success was evaluated by presenting virgin and non-virgin females to the miRNA KO males and calculating the male mating success rate (Lu, et al. 2018). We measured a male’s ability to repress female re-mating according to the re-mating rejection rate of the female (Lu, et al. 2018). Sperm competition was analyzed using by both offense and defense assays (Yeh, et al. 2013; Civetta and Ranz 2019), and we measured male fertility by mating males to two batches of virgin females in series, which aims to exhaust sperm reservoirs in order to obtain an accurate estimate of sperm production (Liufu, et al. 2017).

We compared miR-975 KO in *D. simulans* vs. *D. melanogaster*, which shows no defects in all surveyed components (**Fig. 2B;** also see Lu et al. (2018)). Surprisingly, miR-975 KO in *D. simulans* resulted in severe defects in male mating success (**Fig. 2C**). More than one-third (16 of 43) of mutant *D. simulans* males failed to mate with virgin female, compared with only 1 of 41 wild type males. The mating defect was even more pronounced when miR-975 KO males were presented with non-virgin females, in which the mating success rate drops to only 15% (8 of 53) compared with 73% (32 of 44) in wild type flies. This loss of mating success would substantially decrease fitness due to fewer progeny produced in wild. We also found that miR-975 KO males appeared to have reduced ability to repress female re-mating, but this did not reach statistical significance, perhaps due to the limited numbers tested. According to these data, miR-975 KO seems to have a neutral effect in *D. melanogaster* but was deleterious in *D. simulans*, which echoes its expression difference between the two species (**Fig. 1A**). Compared with the distinct phenotypic effects of miR-975 KO, effects of miR-983 KO were more similar between the two species (**Fig. 2D&E**). Intriguingly, we found that miR-983 KO males had higher offense scores, indicating that the sperm of miR-983 KO males are more competitive. This enhanced performance is offense-specific because we did not observe an elevated score in the defense assay (**Fig. S3**). In addition, we observed that miR-983 KO males likely have a stronger ability to repress female re-mating (**Fig. 2D&E**), and miR-983 KO males produce slightly more progeny in *D. melanogaster* (**Fig. S4**). This observation is surprising because the absence of a highly expressed miRNA might be expected to reduce rather than improves fitness (See further discussion below).

Collectively, these functional assays demonstrate the diverse contributions of the studied miRNAs to male reproduction functions in different species, suggesting the rapid functional evolution of these *de novo* miRNAs.

### Transcriptome divergence between species

The phenotypic consequences of the *de novo* miRNA deletions (miR-983 KO and miR-975 KO) between *D. melanogaster* and *D. simulans* should be a manifestation of the underlying transcriptomic dysregulation. We next compared the number of dysregulated genes in the two species upon miRNA knockout. For miR-975, we found only 396 genes that were significantly dysregulated in *D. melanogaster* (**Fig. 3A**) whereas 1369 genes were dysregulated in *D. simulans* (**Fig. 3B**), a 3-fold difference. This discrepancy, similar to the differences observed in phenotypic assays, might be due to the weak expression of miR-975 in *D. melanogaster* compared to a much higher expression level in *D. simulans* (**Fig. 1A**). Indeed, for miR-983, which has a similar expression in the two species, the magnitude of transcriptomic change was much closer, with 854 genes dysregulated in *D. melanogaster* (**Fig. 3C**) and 1151 in *D. simulans* (**Fig. 3D**). We found the overlap among these mis-regulated genes to be minimal between the two species (**Fig. S5**). Therefore, it is not surprising that dysregulated genes are enriched in distinct gene ontology terms (**Table S2**), indicating significant divergence in target gene regulation between the two species, which results in downstream effects on the entire transcriptome.

**Figure 3.**
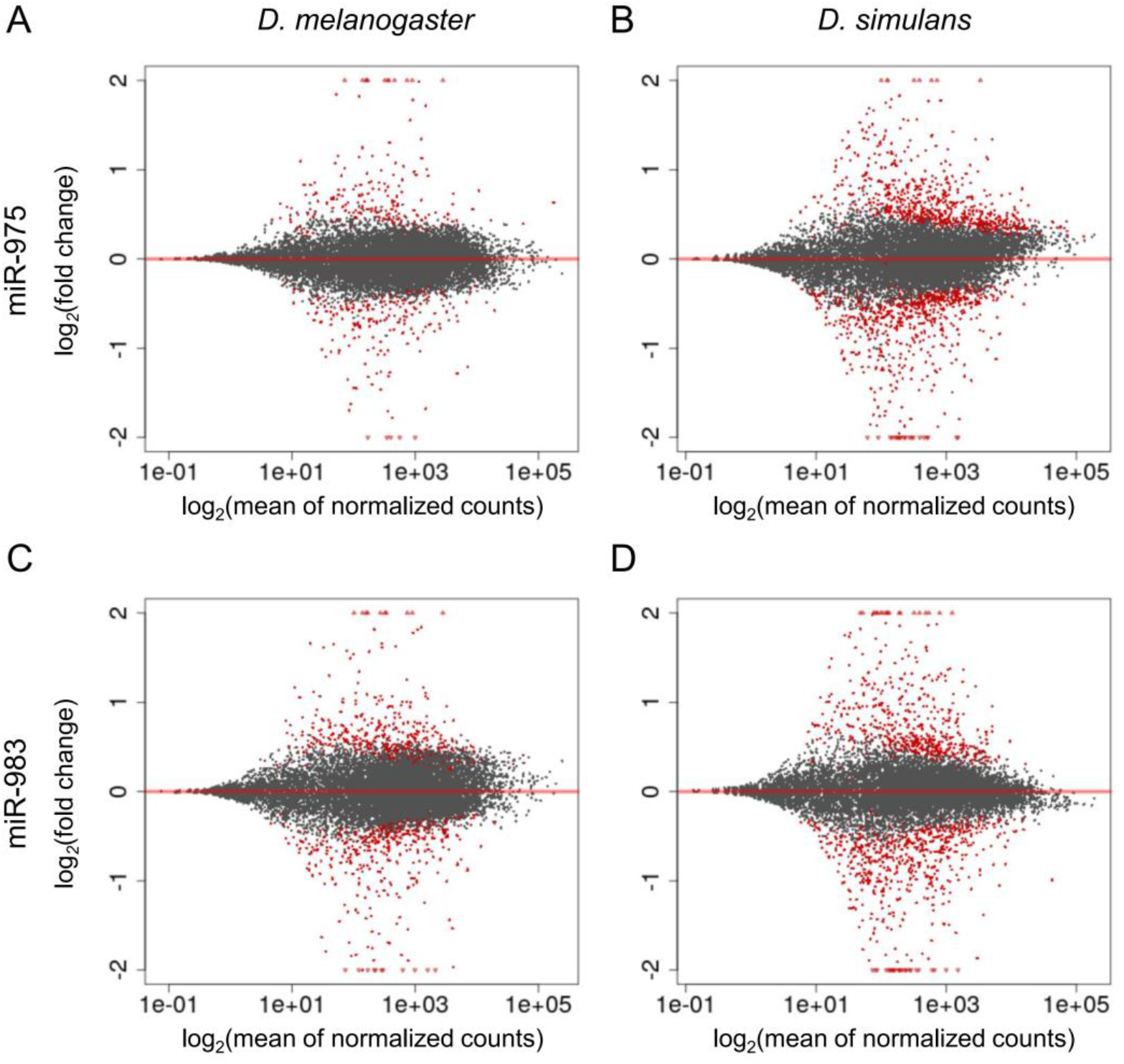
Genome-wide dysregulation in miR-975 and miR-983 KO flies. A) MA plot shows the expression difference between miR-975 KO vs. WT flies in *D. melanogaster*. The significantly differential genes with FDR (false discovery rate) < 0.05 are shown in red dots. B) MA plot shows the expression difference between miR-975 KO vs. WT flies in *D. simulans* C) MA plot shows the expression difference between miR-983 KO vs. WT flies in *D. melanogaster*. D) MA plot shows the expression difference between miR-983 KO vs. WT flies in *D. simulans*.

## Discussion

Van Valen suggested that lower taxa (e.g. genera) are subjected to more unpredictable selective pressure compared to the higher taxa (e.g. classes or orders). Hence, these lower groups often go extinct, even following a period of adaptive existence. In this study, we demonstrated this scenario from a gene-level perspective by focusing on *de novo* miRNA genes.

A unique feature of this study is the choice of genes. The selected *de novo* genes must have been adaptive in the past. Many earlier studies have proposed high turnover rates of new elements in the genome, for both coding (Tautz and Domazet-Loso 2011; Palmieri, et al. 2014) and non-coding genes (Lu, Shen, et al. 2008; Meunier, et al. 2013; Lyu, et al. 2014). As these new elements are mostly non-functional, they may belong to a class often characterized as dead-on-arrival (DOA) elements. The high turnover of neutral DOA elements is unsurprising, whereas new genes that arrive in an adaptive manner are expected to remain functional.

Indeed, our functional assays revealed that miR-975 is functional in *D. simulans*, as deleting the miRNA substantially decreased male mating success, and hence fitness (**Fig. 2C**). However, when the same gene continuously evolves in *D. melanogaster*, a distinct fitness landscape, it may end up in a completely different sequence and functional space due to its neutral contributions to fitness components (**Fig. 2B**). These observations are direct evidence for the Red Queen hypothesis from a gene-level perspective.

In contrast, our assays showed that knockout of miR-983 increased fitness in general for both species. This negative contribution of miR-983 is unexpected given its abundant expression level in both species. In our previous study, we noted that the total fitness effect of miRNAs may be “quasi-neutral”, which means the miRNA KO flies could perform better in some fitness components and worse in others (Lu, et al. 2018). We only measured four components of male reproduction, which may not be sufficient to draw definitive conclusions. However, *de novo* genes die regularly, not only due to complete loss of functionality, but also because of potentially opposing positive and negative fitness effects. This observation indicates that miR-983 may be on an evolutionary death trajectory, as it shows modest negative effects on some fitness components.

The driving force underlying the continual evolution of new genes has been widely discussed (McLysaght and Hurst 2016). Here we should mention that most new genes are specific to male reproduction (Kaessmann 2010; Schlotterer 2015). The arms race in sexual competition is a never-ending battle (Gage 2004; Brennan and Prum 2015; Perry and Rowe 2015; Morimoto, et al. 2019), and selective advantages are often transient. In this sense, sexual selection may very likely operate in a Red Queen landscape. Because new genes, including all the *de novo* miRNAs analyzed here, are often testis-specific in expression, it is plausible that the evolution of new genes may be subject to the Red Queen effect via sexual selection.

## Methods and Materials

Experimental details are included in the supplementary materials. The small RNA-seq libraries generated for this study are curated in GEO with the accession numbers GSM2977400 and GSM2977401. RNA-seq libraries are available in the National Genomics Data Center (https://bigd.big.ac.cn/) with the accession number PRJCA002747.

## Acknowledgements

We thank Prof. Chung-I Wu for his generous assistance and valuable discussion for this study. This work was supported by National Natural Science Foundation of China (31900417) and Guangdong Basic and Applied Basic Research Foundation (2019A1515010708) to G.A.L.

