## supplement for "Run or die in the evolution of new microRNAs - Testing the Red Queen hypothesis on *de novo* new genes"

### Supplementary materials:

**Figure S1 Cross scheme for generating miRNA mutants in *D. simulans***

**Figure S2 Confirmation of miRNA KO by RT-qPCR**

**Figure S3 Defense assay for sperm competition in *D. melanogaster* and *D. simulans***

**Figure S4 Male fertility of miRNA knock out (KO) flies in *D. melanogaster* and *D. simulans***

**Figure S5 Overlap of dysregulated genes between species**

**Table S1 Small RNA libraries**

**Table S2 Gene Ontology analysis for miRNA KO**

**Table S3 TALEN binding sites for generating miRNA KO**

**Table S4 Primers used for PCR**

**Table S5 Fly stocks used in this study**

**Table S6 Primers for RT-qPCR**

**Supplementary Methods and Materials**

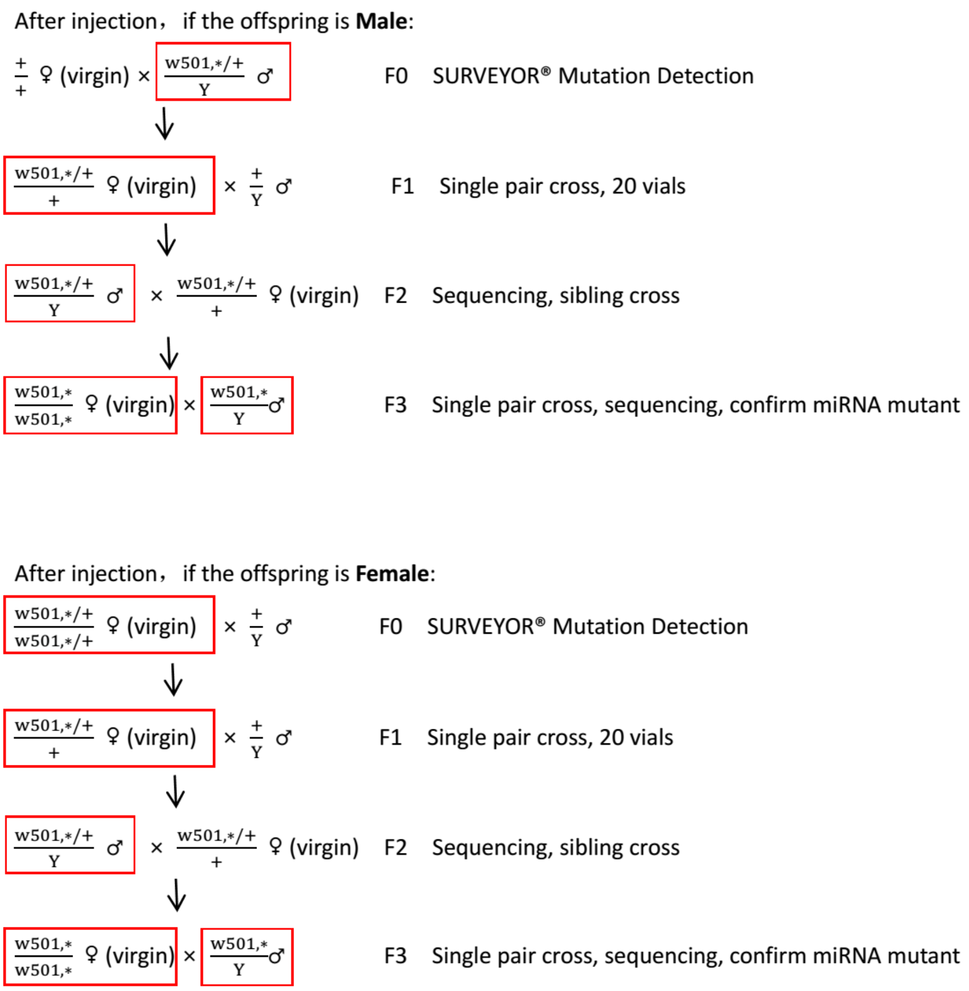


**Figure S1 Cross scheme for generating miRNA mutants in *D. simulans***

Flies in the red boxes are used for mutation detection or Sanger sequencing (methods labeled on the right) after crossing with corresponding females. After embryo injection, each emerging F0 fly was used for mutation detection according to the SURVEYOR® Mutation Detection Kit manufacturer's protocol. If the mutation was detected in F0, twenty F1 virgin females were crossed with wild type red-eye males. F2 males were subjected to Sanger sequencing to detect and confirm the mutant sequences. Flies containing the mutation were crossed with sibling females, making the mutation locus homozygous. F3 flies were subjected to another round of Sanger sequencing to ensure that homozygous mutant flies were chosen. Flies without the mutation served as controls for downstream assays. If F0-emerging flies were female, a similar procedure was used, as depicted in the bottom panel.

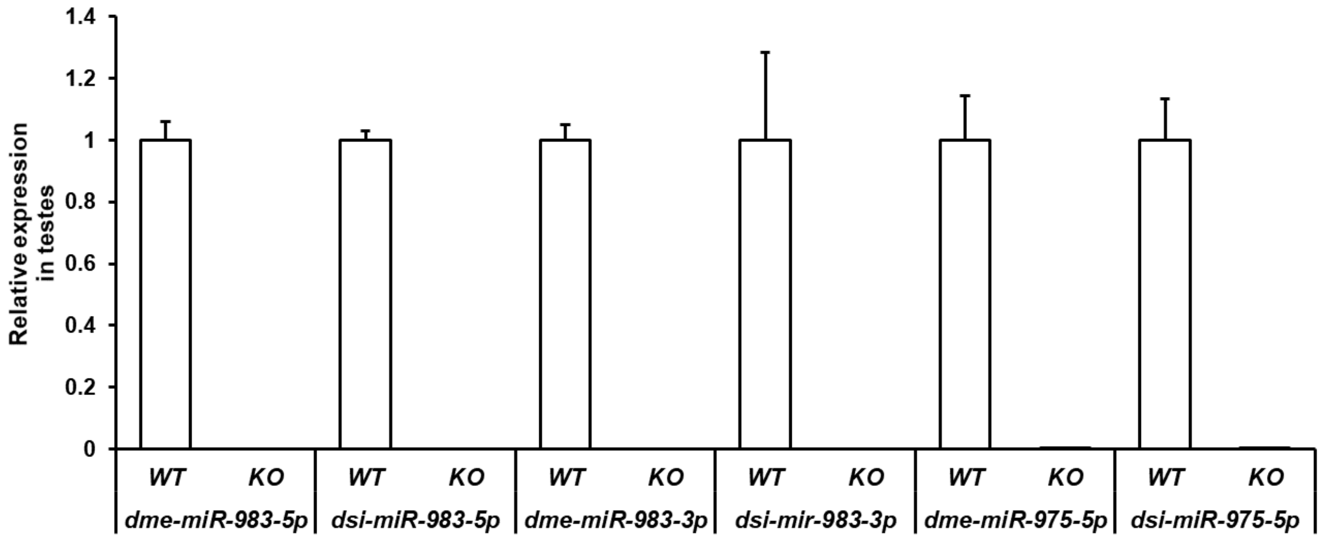


**Figure S2 Confirmation of miRNA KO by RT-qPCR**

*dme-miR-983-5p/-3p*, *dsi-miR-983-5p/-3p, dme-miR-975-5p* and *dsi-miR-975-5p* were undetectable in testes of KO flies. WT, wildtype; KO, knock-out.

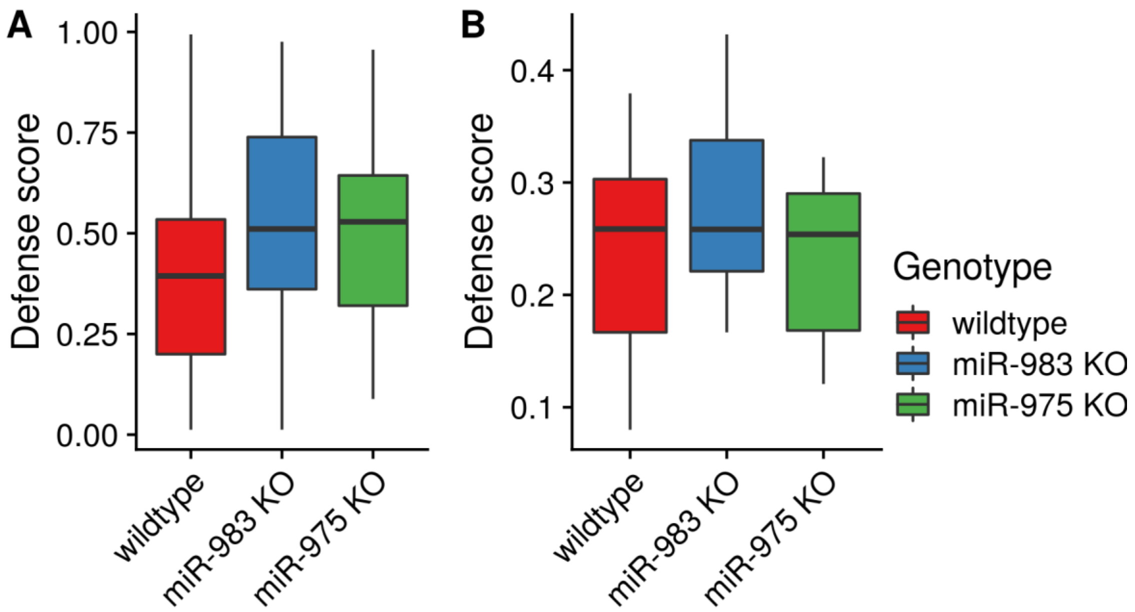


**Figure S3 Defense assay for sperm competition in *D. melanogaster* and *D. simulans***

A) Defense assay in *D. melanogaster.*

B) Defense assay in *D. simulans*. No significant difference was observed.

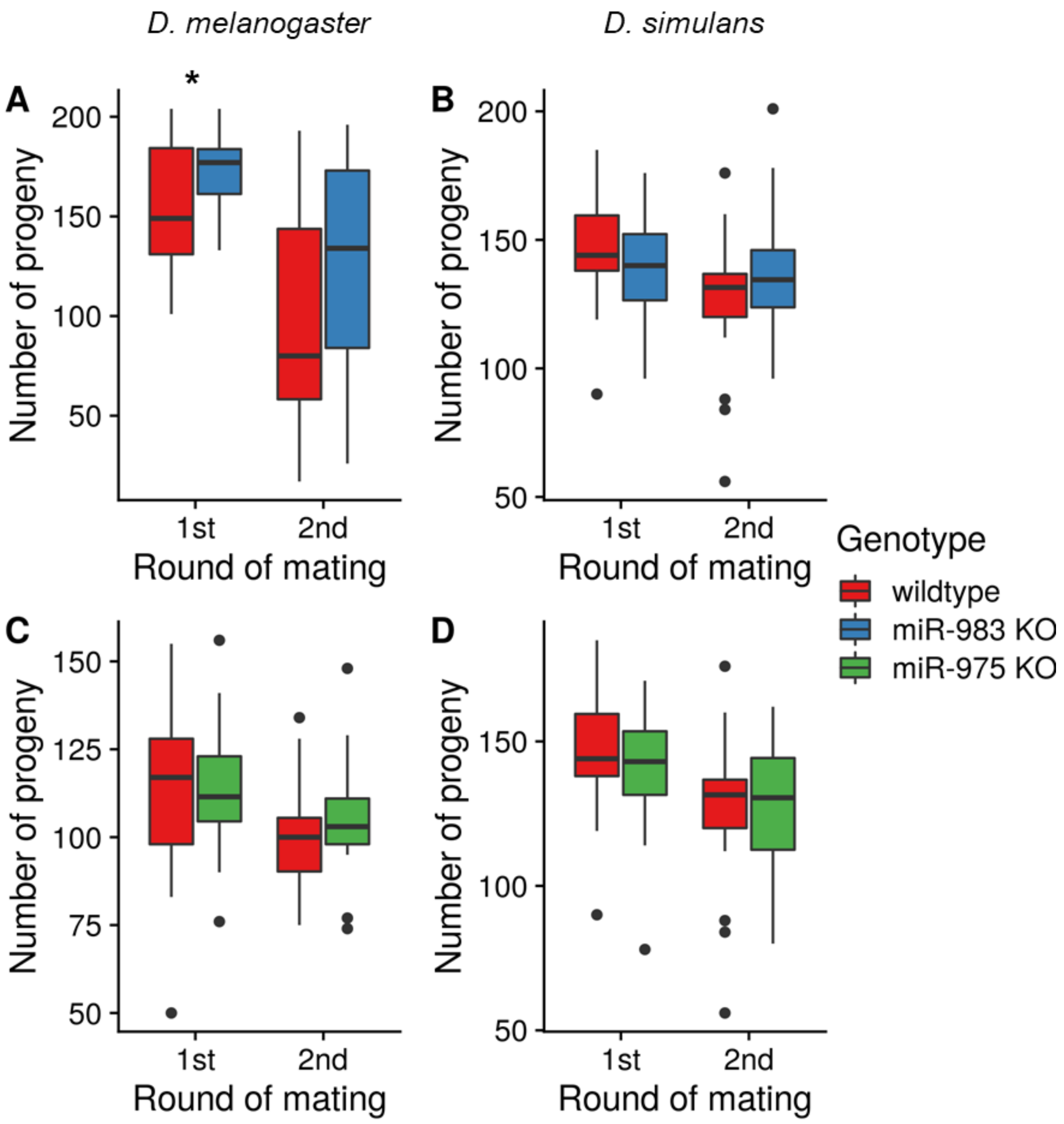


**Figure S4 Male fertility of miRNA knock out (KO) flies in *D. melanogaster* and *D. simulans***

A) Male fertility of miR-983 knock out (KO) flies in *D. melanogaster.*

B) Male fertility of miR-983 KO flies in *D. simulans*.

C) Male fertility of miR-975 KO flies in *D. melanogaster.*

D) Male fertility of miR-975 KO flies in *D. simulans*. All two-sided t-test p-values > 0.05 except the comparison of first-round mating for miR-983 KO vs. wild-type males (p = 0.03).

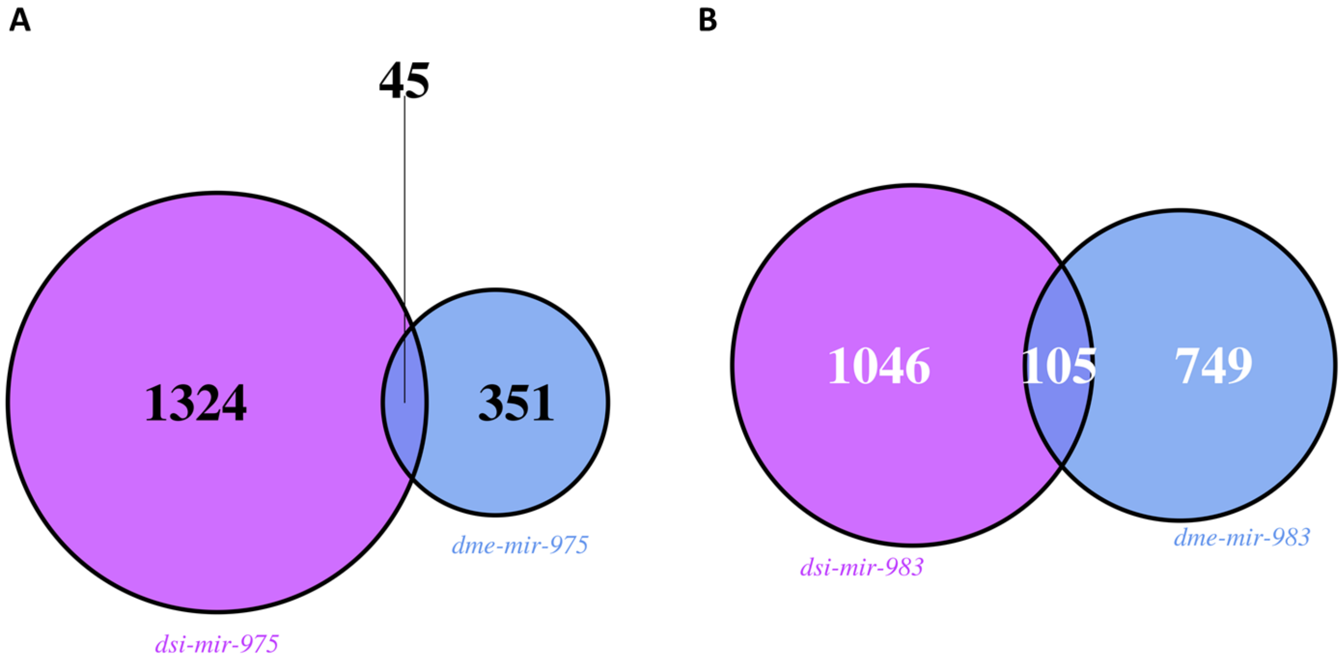


**Figure S5 Overlap of dysregulated genes between species**

A) *dsi-mir-975* KO versus *dme-mir-975* KO.

B) *dsi-mir-983* KO versus *dme-mir-983* KO. Significance is determined by DEseq2 with FDR $< 0.05$.

**Table S1 Small RNA libraries**

| GEO accession | species | citations |
|---|---|---|
| GSM909277 | *D. melanogaster* | (Toledano, et al. 2012) |
| GSM909278 | | |
| GSM280085 | | (Czech, et al. 2008) |
| GSM548582 | | (Rozhkov, et al. 2010) |
| GSM548584 | | |
| GSM548589 | | |
| GSM548591 | | |
| GSM2562978 | | (Zhao, et al. 2018) |
| GSM2562975 | | |
| GSM1165053 | *D. simulans* | (Lyu, et al. 2014) |
| GSM2977400 | *D. sechellia* | this study |
| GSM2977401 | *D. erecta* | |
| GSM548610 | *D. virilis* | (Rozhkov, et al. 2010) |
| GSM548623 | | |

#### Table S2 Gene Ontology analysis for miRNA KO

| species | miRNA | GO.ID | Term | Annotated | Significant | Expected | weight01Fisher |
|---|---|---|---|---|---|---|---|
| *D. mel* | miR-975 | GO:0055085 | transmembrane transport | 543 | 36 | 18.01 | 0.0002 |
| *D. mel* | miR-975 | GO:0017085 | response to insecticide | 22 | 5 | 0.73 | 0.00064 |
| *D. mel* | miR-975 | GO:0042391 | regulation of membrane potential | 64 | 8 | 2.12 | 0.00118 |
| *D. mel* | miR-975 | GO:0006964 | positive regulation of biosynthetic process | 11 | 3 | 0.36 | 0.00489 |
| *D. mel* | miR-975 | GO:0085029 | extracellular matrix assembly | 11 | 3 | 0.36 | 0.00489 |
| *D. mel* | miR-975 | GO:0048100 | wing disc anterior/posterior pattern formation | 11 | 3 | 0.36 | 0.00489 |
| *D. mel* | miR-975 | GO:0030149 | sphingolipid catabolic process | 11 | 3 | 0.36 | 0.00489 |
| *D. mel* | miR-975 | GO:0050877 | nervous system process | 533 | 24 | 17.68 | 0.00615 |
| *D. mel* | miR-975 | GO:0019748 | secondary metabolic process | 93 | 7 | 3.08 | 0.00798 |
| *D. mel* | miR-975 | GO:0034220 | ion transmembrane transport | 259 | 16 | 8.59 | 0.0096 |
| *D. mel* | miR-975 | GO:0019233 | sensory perception of pain | 72 | 7 | 2.39 | 0.00962 |
| *D. mel* | miR-975 | GO:0051260 | protein homooligomerization | 26 | 4 | 0.86 | 0.00997 |
| *D. mel* | miR-975 | GO:0006282 | regulation of DNA repair | 15 | 3 | 0.5 | 0.01223 |
| *D. mel* | miR-975 | GO:0007271 | synaptic transmission, cholinergic | 15 | 3 | 0.5 | 0.01223 |
| *D. mel* | miR-975 | GO:0006672 | ceramide metabolic process | 15 | 3 | 0.5 | 0.01223 |
| *D. mel* | miR-975 | GO:0098754 | detoxification | 26 | 4 | 0.86 | 0.0146 |
| *D. mel* | miR-975 | GO:0016070 | RNA metabolic process | 1619 | 44 | 53.7 | 0.01957 |
| *D. mel* | miR-975 | GO:0006812 | cation transport | 293 | 14 | 9.72 | 0.02323 |
| *D. mel* | miR-975 | GO:0030431 | sleep | 64 | 5 | 2.12 | 0.02516 |
| *D. mel* | miR-975 | GO:0007400 | neuroblast fate determination | 20 | 3 | 0.66 | 0.02712 |

| species | miRNA | GO.ID | Term | Annotated | Significant | Expected | weight01Fisher |
|---|---|---|---|---|---|---|---|
| *D. sim* | miR-975 | GO:0032504 | multicellular organism reproduction | 1147 | 150 | 138.69 | 1.60E-10 |
| *D. sim* | miR-975 | GO:0016485 | protein processing | 61 | 18 | 7.38 | 0.00021 |
| *D. sim* | miR-975 | GO:0010004 | gastrulation involving germ band extension | 37 | 10 | 4.47 | 0.00026 |
| *D. sim* | miR-975 | GO:0016042 | lipid catabolic process | 90 | 11 | 10.88 | 0.00066 |
| *D. sim* | miR-975 | GO:0043455 | regulation of secondary metabolic process | 11 | 6 | 1.33 | 0.00083 |
| *D. sim* | miR-975 | GO:0008380 | RNA splicing | 246 | 27 | 29.75 | 0.00086 |
| *D. sim* | miR-975 | GO:0035006 | melanization defense response | 44 | 8 | 5.32 | 0.001 |
| *D. sim* | miR-975 | GO:0042454 | ribonucleoside catabolic process | 13 | 6 | 1.57 | 0.00176 |
| *D. sim* | miR-975 | GO:0007156 | homophilic cell adhesion via plasma membrane | 40 | 12 | 4.84 | 0.00202 |
| *D. sim* | miR-975 | GO:0018401 | peptidyl-proline hydroxylation to 4-hydroxy | 21 | 8 | 2.54 | 0.00212 |
| *D. sim* | miR-975 | GO:0051049 | regulation of transport | 226 | 23 | 27.33 | 0.00653 |
| *D. sim* | miR-975 | GO:0007608 | sensory perception of smell | 79 | 18 | 9.55 | 0.00769 |
| *D. sim* | miR-975 | GO:0035160 | maintenance of epithelial integrity | 12 | 5 | 1.45 | 0.00976 |
| *D. sim* | miR-975 | GO:0002027 | regulation of heart rate | 12 | 5 | 1.45 | 0.00976 |
| *D. sim* | miR-975 | GO:0035289 | posterior head segmentation | 13 | 5 | 1.57 | 0.01431 |
| *D. sim* | miR-975 | GO:0000578 | embryonic axis specification | 113 | 11 | 13.66 | 0.01479 |
| *D. sim* | miR-975 | GO:0006518 | peptide metabolic process | 498 | 66 | 60.22 | 0.01568 |
| *D. sim* | miR-975 | GO:0045465 | R8 cell differentiation | 20 | 5 | 2.42 | 0.01624 |
| *D. sim* | miR-975 | GO:0046907 | intracellular transport | 629 | 57 | 76.06 | 0.01744 |
| *D. sim* | miR-975 | GO:0032880 | regulation of protein localization | 87 | 10 | 10.52 | 0.02036 |

#### Table S2 (Continued)

| species | miRNA | GO.ID | Term | Annotated | Significant | Expected | weight01Fisher |
|---|---|---|---|---|---|---|---|
| *D. mel* | miR-983 | GO:0055085 | transmembrane transport | 532 | 52 | 39.97 | 0.0012 |
| *D. mel* | miR-983 | GO:0006030 | chitin metabolic process | 73 | 13 | 5.48 | 0.0028 |
| *D. mel* | miR-983 | GO:1901568 | fatty acid derivative metabolic process | 25 | 4 | 1.88 | 0.0056 |
| *D. mel* | miR-983 | GO:0009914 | hormone transport | 17 | 3 | 1.28 | 0.0056 |
| *D. mel* | miR-983 | GO:0097194 | execution phase of apoptosis | 29 | 5 | 2.18 | 0.0096 |
| *D. mel* | miR-983 | GO:0042737 | drug catabolic process | 51 | 9 | 3.83 | 0.0127 |
| *D. mel* | miR-983 | GO:0009620 | response to fungus | 55 | 11 | 4.13 | 0.0128 |
| *D. mel* | miR-983 | GO:0018401 | peptidyl-proline hydroxylation to 4-hydroxy | 20 | 5 | 1.5 | 0.0142 |
| *D. mel* | miR-983 | GO:0055114 | oxidation-reduction process | 511 | 44 | 38.39 | 0.0157 |
| *D. mel* | miR-983 | GO:0006508 | proteolysis | 735 | 58 | 55.22 | 0.0171 |
| *D. mel* | miR-983 | GO:0015696 | ammonium transport | 21 | 5 | 1.58 | 0.0175 |
| *D. mel* | miR-983 | GO:0009584 | detection of visible light | 31 | 5 | 2.33 | 0.0177 |
| *D. mel* | miR-983 | GO:0035249 | synaptic transmission, glutamatergic | 22 | 5 | 1.65 | 0.0213 |
| *D. mel* | miR-983 | GO:0007628 | adult walking behavior | 15 | 4 | 1.13 | 0.0221 |
| *D. mel* | miR-983 | GO:0007472 | wing disc morphogenesis | 261 | 23 | 19.61 | 0.0251 |
| *D. mel* | miR-983 | GO:0048580 | regulation of post-embryonic development | 20 | 4 | 1.5 | 0.0252 |
| *D. mel* | miR-983 | GO:0062013 | positive regulation of small molecule metabolic process | 16 | 4 | 1.2 | 0.0278 |
| *D. mel* | miR-983 | GO:0016239 | positive regulation of macroautophagy | 16 | 4 | 1.2 | 0.0278 |
| *D. mel* | miR-983 | GO:0042073 | intraciliary transport | 16 | 4 | 1.2 | 0.0278 |
| *D. mel* | miR-983 | GO:0042332 | gravitaxis | 24 | 5 | 1.8 | 0.0303 |

| species | miRNA | GO.ID | Term | Annotated | Significant | Expected | weight01Fisher |
|---|---|---|---|---|---|---|---|
| *D. sim* | miR-983 | GO:0032504 | multicellular organism reproduction | 1147 | 121 | 115.99 | 1.50E-09 |
| *D. sim* | miR-983 | GO:0018401 | peptidyl-proline hydroxylation to 4-hydroxy | 21 | 10 | 2.12 | 1.30E-05 |
| *D. sim* | miR-983 | GO:0055114 | oxidation-reduction process | 515 | 68 | 52.08 | 1.50E-05 |
| *D. sim* | miR-983 | GO:0070192 | chromosome organization involved in meiotic cell cycle | 47 | 11 | 4.75 | 0.0018 |
| *D. sim* | miR-983 | GO:0019359 | nicotinamide nucleotide biosynthetic process | 33 | 5 | 3.34 | 0.0028 |
| *D. sim* | miR-983 | GO:0010951 | negative regulation of endopeptidase activity | 34 | 9 | 3.44 | 0.0054 |
| *D. sim* | miR-983 | GO:0007606 | sensory perception of chemical stimulus | 166 | 29 | 16.79 | 0.0093 |
| *D. sim* | miR-983 | GO:0044257 | cellular protein catabolic process | 302 | 27 | 30.54 | 0.0103 |
| *D. sim* | miR-983 | GO:0007135 | meiosis II | 15 | 5 | 1.52 | 0.0132 |
| *D. sim* | miR-983 | GO:0070873 | regulation of glycogen metabolic process | 10 | 4 | 1.01 | 0.0133 |
| *D. sim* | miR-983 | GO:0010906 | regulation of glucose metabolic process | 16 | 7 | 1.62 | 0.0162 |
| *D. sim* | miR-983 | GO:0072528 | pyrimidine-containing compound biosynthetic process | 23 | 5 | 2.33 | 0.0163 |
| *D. sim* | miR-983 | GO:1901658 | glycosyl compound catabolic process | 19 | 4 | 1.92 | 0.0163 |
| *D. sim* | miR-983 | GO:0030100 | regulation of endocytosis | 53 | 7 | 5.36 | 0.0178 |
| *D. sim* | miR-983 | GO:0007608 | sensory perception of smell | 79 | 15 | 7.99 | 0.0189 |
| *D. sim* | miR-983 | GO:0045143 | homologous chromosome segregation | 11 | 4 | 1.11 | 0.0192 |
| *D. sim* | miR-983 | GO:0006575 | cellular modified amino acid metabolic process | 90 | 18 | 9.1 | 0.0225 |
| *D. sim* | miR-983 | GO:0006633 | fatty acid biosynthetic process | 57 | 10 | 5.76 | 0.0247 |
| *D. sim* | miR-983 | GO:0050790 | regulation of catalytic activity | 305 | 32 | 30.84 | 0.0265 |
| *D. sim* | miR-983 | GO:0033617 | mitochondrial respiratory chain complex | 12 | 4 | 1.21 | 0.0265 |

**Table S3 TALEN binding sites for generating miRNA KO**

| miRNA | TALEN binding site |
|---|---|
| ***dsi-mir-975*** | TTTTGATTTTAAACACTGCCTACATCCTGTATGTGTTTTGCATCCG |
| ***dsi-mir-983*** | GACTGAAAATCATGTGTTCGAAAGGTATTACTAAATTACTGCAAG |

Mature miRNA sequences are in red, and the left and right TALEN binding sites are underlined. Note that the reverse complement sequence for *dsi-mir-983* is shown here.

**Table S4 Primers used for PCR**

| miRNA | F primer | R primer | Fragment Length |
|---|---|---|---|
| ***dsi-mir-975*** | TCGGCACTGGAGTGCTAAAT | GCTTCGTCACTACGTGGACA | 820 |
| ***dsi-mir-983*** | GCACGGTACGTTTATGTTTCTAGG | GGTAAAGAAGGGAAAGCTTTTGGG | 899 |

**Table S5 Fly stocks used in this study**

| Species | Genotype |
|---|---|
| ***D. simulans*** | $w^{501}$ |
| ***D. simulans*** | *Sim6* |
| ***D. erecta*** | wildtype (Stock Number: 14021-0224.01) |
| ***D. sechellia*** | wildtype (Stock Number: 14021-0248.20) |

**Table S6 Primers for RT-qPCR**

Reverse transcription primers

| miRNA | RT Primer |
|---|---|
| ***dme-miR-983-5p*** | GTTGGCTCTGGTGCAGGGTCCGAGGTATTCGCACCAGAGCCAAC**TCATTA** |
| ***dme-miR-983-3p*** | GTTGGCTCTGGTGCAGGGTCCGAGGTATTCGCACCAGAGCCAAC**AGATAA** |
| ***dsi-miR-983-5p*** | GTTGGCTCTGGTGCAGGGTCCGAGGTATTCGCACCAGAGCCAAC**TCATGT** |
| ***dsi-miR-983-3p*** | GTTGGCTCTGGTGCAGGGTCCGAGGTATTCGCACCAGAGCCAAC**TAGATA** |
| ***dme-miR-975-5p*** | GTTGGCTCTGGTGCAGGGTCCGAGGTATTCGCACCAGAGCCAAC**ATACAG** |
| ***dsi-miR-975-5p*** | GTTGGCTCTGGTGCAGGGTCCGAGGTATTCGCACCAGAGCCAAC**ATACAG** |
| **2s** | GTTGGCTCTGGTGCAGGGTCCGAGGTATTCGCACCAGAGCCAACTACAAC |

Green regions show the reverse complement sequences for the last 6 nt of the miRNA 3' end.

Quantitative real-time PCR primers

| miRNA | Primes |
|---|---|
| ***dme-miR-983-5p*** | GGCG**ATAATACGTTTCGAAC** |
| ***dme-miR-983-3p*** | GGCG**ATTAGGTAGTTACGCA** |
| ***dsi-miR-983-5p*** | GGCG**AGTAATACCTTTCGAAC** |
| ***dsi-miR-983-3p*** | GGCG**ATGAGTTCGTCAGGTAT** |
| ***dme-miR-975-5p*** | GGCG**TAAACACTTCCTACATC** |
| ***dsi-miR-975-5p*** | GGCG**TAAACACTGCCTACATC** |
| **2s** | GGCGTGCTTGGACTACATATGG |
| **Universal reverse primer** | GTGCAGGGTCCGAGGT |

Red regions show the mature miRNA sequences (not including the last 6 nt at 3' end).

**Methods and Materials:**

*Small RNA analyses*

Testes were dissected and collected from 3-5-day-old *D. erecta* (UCSD stock #: 14021-0224.01) and *D. sechellia* (stock #: 14021-0248.20) adults. Total RNA was extracted using the TRIzol® Reagent. Small RNA libraries were generated using an Illumina Small RNA Sample Preparation kit and subsequently sequenced using an Illumina HiSeq 2000 machine at the Beijing Genomics Institute (Shenzhen).

To determine the expression patterns of *de novo* miRNAs, we also collected and surveyed nine testes libraries in *D. melanogaster*, one in *D. simulans,* and two in *D. virilis*. Sequencing data were retrieved from the GEO database (http://www.ncbi.nlm.nih.gov/geo/), and accession numbers are shown in **Table S1** (Edgar et al. 2002). Fly genomes were retrieved from FlyBase (http://flybase.org/) (Attrill et al. 2016). The genome versions used were: *D. melanogaster*: r6.04, *D. simulans*: r2.01, *D. sechellia*: r1.3, *D. erecta*: r1.04, and *D. virilis*: r1.03. miRNA precursor orthologous sequences were adopted from Mohammed et al. (Mohammed, et al. 2013). Mature miRNA sequences were retrieved from miRBase (http://www.mirbase.org; release 21) (Kozomara and Griffiths-Jones 2014). miRNA expression was measured using the Mapper and Quantifier modules in miRDeep2 (version: 2.0.0.7). Short reads were mapped to genomes using mapper.pl with default parameters, allowing no mismatches in the first 18 nt of reads and no more than two mismatches in the remainder of the reads. Reads matching miRNAs in each library were normalized using all reads mappable to mature sequences and scaled as Reads Per Million (RPM).

*miRNA mutant fly construction and stocks*

miRNA mutant flies were generated using a TALEN (Transcription Activator-Like Effector Nuclease) method (Katsuyama, et al. 2013). TALEN plasmids were designed and constructed by ViewSolid Biotech Co., Ltd (http://www.v-solid.com/). TALEN binding sites are shown in supplementary **Table S3**. Plasmids were transcribed to mRNA *in vitro* using the mMESSAGE mMACHINE® T7 ULTRA Transcription Kit. $w^{501}$ embryos were used for microinjection in *D. simulans*. After microinjection, embryos were kept at 25°C, and emerging adult flies were crossed to the *Sim6* strain in *D. simulans* (see **Fig. S1** for a detailed cross scheme). DNA was extracted from $F_0$ adult flies after five days of crossing. PCR was performed to amplify the regions containing miR-983 and miR-975 in *D. simulans* for mutation detection using the Surveyor® Mutation Detection Kit. Once a mutation was detected, single-pair crossing was performed with $F_1$ flies. Sanger sequencing of mutants was performed to determine InDel sequences. Primers used for PCR are shown in **Table S4**. Flies without miRNA mutations were subjected to the same cross scheme and used as controls in subsequent assays. All procedures using commercial kits were performed following manufacturers' protocols. Fly stocks used in this study are listed in **Table S5**.

*Quantitative miRNA analyses by qRT-PCR*

We measured relative expression levels of miRNAs in the miRNA knock out and control strains. Total RNA was extracted from testes of 3-5-day-old flies using the TRIzol® Reagent (Thermo Fisher Scientific Inc., catalog # 15596026). We used stem-loop primers for reverse transcription (Chen, et al. 2005), and Taqman PCR analysis was then performed using the miRNA UPL (Roche Diagnostics) probe assay protocol (Varkonyi-Gasic, et al. 2007; He, et al. 2016). Primers are listed in **Table S6**. Three biological replicates were conducted for each genotype, and *2s* RNA was used as an endogenous control.

*Phenotypic assays*

*Male fertility*

Two rounds of mating were performed with miR-983 KO and miR-975 KO flies. In *D. melanogaster*, each 3-5-day-old male was mated with five 3-5-day-old $w^{1118}$ virgin females for two days, and then transferred to a new vial to mate with another five virgin females for two days. In *D. simulans*, the two mating procedure rounds were the same, except that *D. simulans* control virgin females were used for mating. All offspring were counted in each of the two consecutive mating rounds. More than 15 males were examined for each genotype in *D. melanogaster* and *D. simulans.* Male fertility was measured as the average number of offspring per male. Two-tailed t-test was used to determine the statistical significance of phenotypic differences between genotypes.

*Mating success and Male ability to repress female re-mating*

Three distinct metrics were measured: male mating success with virgin females, male mating success with pre-mated females, and males' ability to repress female re-mating. To survey mating success with virgin females, wild-type virgin females were mated to either wild-type or miRNA knockout males for 2h. Mating success rates for the males were inferred by whether progeny were produced. This was calculated according to the number of females with progeny divided by the total number of females used in the assay. To survey male mating success with non-virgin females, wild-type females were first mated to reference males for 2h. After 2.5 d, these mated females were presented with either wild-type or miRNA knockout males for 12h. Male mating success with non-virgin females was calculated by the number of females with both white-eyed and red-eyed progeny (successful fertilization by experimental and reference males, respectively) divided by the total number of females successfully pre-mated with reference males. To survey males' ability to repress female re-mating, virgin females were mated to either wild-type or miRNA knockout males for 2h. After 2.5 d, these mated females were presented with reference males for 12h. We measured two outcomes: 1) number of females (x) with white-eyed progeny only (rejection of the reference male), and 2) number of females (y) with both white-eyed and red-eyed progeny (acceptance of the reference male). Then, male ability to repress female re-mating was calculated as x / (x+y). The reference males were from the *Sim6* line in *D. simulans*. P-values were calculated using Fisher's exact test. *D. melanogaster* data from Lu et al. (2018) were re-used for analysis.

*Sperm competition*

Sperm competition assays were conducted as described in Yeh, et al. (2013). Briefly, double-mating experiments were set up for each miRNA KO and wild-type line. In *D. simulans*, virgin wild-type females were mated to reference males (a wild type red eye strain, *Sim6*) for 2h and then mated to experimental males (miRNA KO or wild-type) for 12h after 56h to measure offensive ability. Conversely, wild-type females were mated to experimental males for 2h and then to the reference males for 12h after 56h for the defense assay.

Each assay used 40-50 replicates. Progeny eye color was used as a marker of mating success and paternity identification. $F_0$ females with both red-eyed and white-eyed progeny had successfully mated with both experimental and reference males. Red-eyed female progeny (y) arose from reference males, whereas white-eyed female progeny (x) were generated from experimental fathers. Sperm competitive ability was estimated using the score $P = x / (x+y)$. Mann-Whitney *U* test was used to determine the statistical significance of phenotypic differences between genotypes. *D. melanogaster* data from Lu, et al. (2018) were re-used for analysis.

*Transcriptome analyses*

To detect the effects of miR-983 and miR-975 KO on the transcriptome, we performed RNA-seq analysis. Total RNA was extracted from ~30 testes of 3-5-day-old males using the TRIzol® Reagent. PolyA-enriched RNA libraries were constructed and sequenced on an Illumina HiSeq 2000 at GENEWIZ Co. (https://www.genewiz.com/). One *dme-mir-975* KO sample with an abnormal GC content (42%) was removed and not used for further analysis. *D. melanogaster* (r6.30) and *D. simulans* (r2.02) genomes were downloaded from FlyBase and used for mapping. Raw RNA-seq data were mapped with default parameters using HISAT2 (Pertea, et al. 2016) and then quantified with StringTie (Pertea, et al. 2015). Genes with 1:1 orthologs for *D. melanogaster* and *D. simulans* were used for analysis. DEseq2 was used to detect significantly dysregulated genes with $FDR < 0.05$ (Love, et al. 2014). Gene Ontology analysis was performed using the R package topGO (Alexa, et al. 2020). RNA-seq libraries are available in the National Genomics Data Center (https://bigd.big.ac.cn/) with the accession number PRJCA002747.
